# Integrative Pharmacogenomics Analysis of Patient Derived Xenografts

**DOI:** 10.1101/471227

**Authors:** Arvind Singh Mer, Wail Ba-alawi, Petr Smirnov, Yi Xiao Wang, Ben Brew, Janosch Ortmann, Ming-Sound Tsao, David Cescon, Anna Goldenberg, Benjamin Haibe-Kains

## Abstract

One of the key challenges in cancer precision medicine is finding robust biomarkers of drug response. Patient-derived tumor xenografts (PDXs) have emerged as reliable preclinical models since they better recapitulate tumor response to chemo- and targeted therapies. However, the lack of standard tools poses a challenge in the analysis of PDXs with molecular and pharmacological profiles. Efficient storage, access and analysis is key to the realization of the full potential of PDX pharmacogenomic data. We have developed Xeva (XEnograft Visualization & Analysis), an open-source software package for processing, visualization and integrative analysis of a compendium of *in vivo* pharmacogenomic datasets. The Xeva package follows the PDX minimum information (PDX-MI) standards and can handle both replicate-based and 1×1×1 experimental designs. We used Xeva to characterize the variability of gene expression and pathway activity across passages. We found that only a few genes and pathways have passage specific alterations (median intraclass correlation of 0.53 for genes and positive enrichment score for 92.5% pathways). For example, activity of the mRNA 3’-end processing and elongation arrest and recovery pathways were strongly affected by model passaging (gene set enrichment analysis false discovery rate [FDR] <5%). We then leveraged our platform to link the drug response and the pathways whose activity is consistent across passages by mining the Novartis PDX Encyclopedia (PDXE) data containing 1,075 PDXs spanning 5 tissue types and 62 anticancer drugs. We identified 87 pathways significantly associated with response to 51 drugs (FDR < 5%), including associations such as erlotinib response and signaling by EGFR in cancer pathways and MAP kinase activation in TLR cascade and binimetinib response. Among the significant pathway-drug associations, we found novel biomarkers based on gene expressions, Copy Number Aberrations (CNAs) and mutations predictive of drug response (concordance index > 0.60; FDR < 0.05). Xeva provides a flexible platform for integrative analysis of preclinical *in vivo* pharmacogenomics data to identify biomarkers predictive of drug response, a major step toward precision oncology.

## INTRODUCTION

Preclinical models are vital for investigating disease biology and therapeutics, constituting essential tools for translational research and drug development. In cancer research, immortalized cell lines are the most used preclinical models because of their low cost, flexibility and the existence of assays enabling genetic and chemical screen in a high-throughput manner. These *in vitro* pharmacogenomic studies led to the discovery of many clinically-approved biomarkers for anticancer therapies^1^. Recent large-scale *in vitro* drug screening datasets^2–4^, coupled with rigorous computational analysis pipelines^5–11^ hold the promise to find new drug response biomarkers. However, the cell line models suffer from multiple limitations. Although they are derived from patient tumors, they have evolved to survive in artificial culture conditions resulting in major alterations at the genomic level^12–17^. These *in vitro* models also lack the tumor heterogeneity and three-dimensional structure of the origin patient tumor^15,18–20^.

To create cancer models that better recapitulate the tumor molecular features and drug response, the pharmaceutical industry and academia massively invested in the development of patient-derived xenografts (PDXs)^21^, which enable engraftment of human tumors in animal models^22–28^. PDXs are created by subcutaneous or orthotopic engraftment of the cancerous tissues or cells from patients’ tumors into immunodeficient mice. Once established, these tumors can be passed from mouse to mouse, leading to consecutive “passages” of the initial tumor cells.

PDX models are being generated and distributed by several academic groups, research institutes and commercial organizations. This makes it challenging to find PDX models with specific characteristics such as a model with a specific mutation. Therefore, catalogues of PDX models (e.g. PDXFinder^29^, EurOPDX^30^ and PRoXe^31^) are being developed which contain relevant information and provide links to model acquisition. Furthermore, as PDXs are becoming the gold standard model for preclinical studies, better data standardization and analysis platforms are required to ensure consistency and reproducibility in PDX-based analysis. Recently, a robust standard called PDX models minimal information (PDX-MI) has been proposed^32^ for reporting and quality assurance for PDX models. However, management, analysis and visualization of the PDX-based drug screening and genomic data still constitute major challenges.

Here we present Xeva (Xenograft Visualization and Analysis), a computational package enabling storage, access and analysis of *in vivo* pharmacogenomics data. The Xeva toolbox facilitates biomarker discovery in PDX-based pharmacogenomic data. It implements class structure to manage and connect PDX-based drug screening data to the genomic features of the corresponding tumor. It provides functions for PDX data analysis, including multiple metrics to summarize drug response for tumor growth curves. Xeva provides functions to compute the association between genomic features and response to a drug in PDXs (gene-drug association). Different response metrics for the PDX growth curve can be used as outcome to identify novel gene-drug associations that can act as drug companion tests. Using the Xeva platform, we analyzed gene expression of PDXs across passages and demonstrated that activity patterns of the majority of the genes and pathways are stable across different PDX passages. Our analysis shows that PDXs maintain the vast majority of the pathway activity across passages. We identified multiple pathways significantly associated with anticancer drug response, including known and new biomarkers based on gene expression, copy number aberrations (CNAs) and mutations. Our results support the value of large-scale PDX-based drug screening for biomarker discovery using the integrative pharmacogenomic analysis pipelines implemented in the Xeva computational platform.

## MATERIALS AND METHODS

### Processing of Pharmacogenomic Data

Gene expression data, PDX passages and tissue information were obtained from the Gene Expression Omnibus series GSE78806. Gene expression profiles were normalized with the RMA algorithm in the *Affy* package (version 1.58.0) in Bioconductor^33^. PDX passages and tissue information were curated manually. Molecular profiles including mutation, CNA, gene expression and pharmacological profiles were obtained from the publication Gao *et al.*^28^. Data were processed using R statistical software (http://www.r-proiect.org/). The final dataset contains 3,470 unique PDX models tested across 57 treatments and derived from 191 patients spanning across 5 different cancer types.

### Implementation

PDX pharmacogenomic experiments aim to investigate how tumor volume changes in *in vivo* models with respect to time, with and without drug treatment. The corresponding metadata, such as drug dose, number of days from tumor implantation to treatment start or reason for stopping the experiment (e.g. whether mice died because of complications or were sacrificed due to maximal allowed tumor volume reached), are crucial factors for downstream analysis. In the Xeva platform, we have developed the *XevaSet* class to effectively store pharmacological response (time vs. tumor volume) of PDXs along with metadata related to the experiments and molecular data. Furthermore, to store individual PDX (mouse) model data, we have implemented the *pdxModel* class, which provides slots for PDX-MI variables, along with time vs. tumor volume data. Detailed schematics of the *XevaSet* and *pdxModel* classes are shown in Supplementary Figure S2 and S3, respectively. *XevaSet* object can contain multimodal genomic data, which are linked to individual xenograft models and their pharmacological profiles. Data can be subsetted by metadata and experimental factors, such as tumor types or drug names.

### PDX Experiment Design

*In vivo* evaluation of drug sensitivity in cancer xenografts has traditionally used experimental approaches incorporating multiple animals (five to ten) replicates for each control and treatment arm. This experimental design allows the assessment of the variability in the PDX drug response across individual animals^34^ However, scaling this strategy for high-throughput drug screening is costly and requires the use of a large number of experimental animals. To increase the number of compounds being tested, reduce cost and permit the use of fewer animals to provide essential data^35^, a simpler “1×1×1” experimental design for PDX clinical trial (PCT) has been proposed^28, 36^. In this design, each compound is tested in only one PDX model from each patient (Supplementary Figure S9). Though the 1×1×1 experiment design allows high-throughput screening at reduced cost, it is more prone to experimental errors due to lack of replications. Factors such as difference in the handling of mice or measurement errors in the tumor volume and biological variability in tumor growth rates between animals could significantly impact the results. We foresee that both experimental design strategies will co-exist, where the 1×1×1 experiment design will be used for population-level and high-throughput screening, while the replicate-based experiment design will be used for more focused studies where discrimination of small differences in drug response is a desired outcome. We therefore implemented functions to accommodate both experimental designs for data visualization and statistical analyses.

### PDX Response Metrics

Given the lack of consensus regarding the best summary metrics to estimate *in vivo* drug response^28,37–40^, we implemented the state-of-the-art response metrics used for PDX-based drug response experiments. These include the slope of curves, angle between the mean control and treatment curves, tumor growth inhibition (TGI), area between the curves, linear mixed model^39^, best average response (BAR), best response (BR) and mRECIST^28^.

For each PDX model, least squares fits were obtained by regressing tumor volume at each time point as:

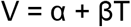

where V denotes tumor volume, T denotes time. The intercept and slope are denoted by α and β, respectively. Subsequently, the angle was computed using inverse tangent of regression line slope as:

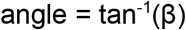

The tumor growth inhibition (TGI) is defined as:

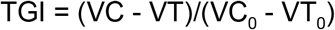

where VC and VT are the median of control and treated growth curve respectively at the end of the study. VC_0_ and VT_0_ indicate the initial tumor volume for control and treated growth curve respectively.

The best average response (BAR) metric for each PDX model is defined as follows:

At each time point t, the normalized change in tumor volume is computed as

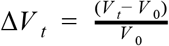

Next, for each time t, the running average of Δ*V_t_* from *t*=0 to *t*, was calculated. BAR is defined as the minimum of this running average (at *t* ≥10 *days*). The minimum value of Δ*V* (at *t*≥10 *days*) defines as best response (BR). Using these, the mRESIST metric for PDX is computed as:

- complete response (CR): BR < −95% and BAR < −40%
- partial response (PR): BR < −50% and BAR < −20%
- stable disease (SD): BR < 35% and BAR < 30%
- progressive disease (PD): not otherwise categorized

### Visualization of *In Vivo* drug Screening Data

Several Xeva functions enable multi-faceted visualization and exploration of the PDX data. The *plotPDX* function displays tumor growth curves, plotting time versus tumor volume data for a patient-drug pair or matched control and treatment PDX models (called batch). The *waterfall* function visualizes population-level response for a given set of PDXs. Similarly, *plotmRECIST* displays mRECIST-based drug response as a heatmap, with drugs along heatmap rows and PDXs along columns. Heatmap cells are colored according to the mRECIST value for the corresponding PDX-drug pair.

### Gene Expression Consistency Analysis

We calculated the Pearson correlation coefficient between pairs of samples belonging to the same lineage using the L1000 landmark gene set^41^. For comparison, we also computed the Pearson correlation coefficient between all possible pairs of samples that do not belong to the same lineage. For genes, we computed the intra-class correlation coefficient (ICC) using the psych package (version 1.8.4). For a particular gene, in each sample we computed its rank by sorting the expression values. The rank of genes along with passage information is used for ICC calculation. ICC values of the genes were used to perform gene set enrichment analysis on the MSigDB hallmark gene sets^42^ and Reactome pathway database^43^, which contains gene sets associated with specific pathways.

### Pathways and Gene-Drug Association Analysis

We computed the association between a molecular feature and response to a drug across PDXs (commonly referred to as gene-drug association or drug response predicting biomarker). The gene-drug association was assessed separately for all three available molecular features, i.e. gene expression, CNA and gene mutation. The response of PDXs to a drug treatment is defined using best average response (BAR).

The association between genomic feature and PDX response was computed using non-parametric measure of association, *concordance index (CI)*^44^ and its equivalent *Somers’ Dxy rank correlation (DXY)*^45^. The CI represents the probability that two variables will rank a random pair of samples the same order. The DXY is equivalent to rescaling the CI values between −1 and 1 using (CI – 0.5) * 2.

For each drug, we compiled a list of potential biomarkers using OncoKB^46^ and literature curation. We adjusted the p-value using FDR for drug combinations and drugs with multiple potential biomarkers. Univariate gene-drug associations were calculated for three different molecular profiling modalities that are gene expression, copy number variation and mutation. For drug-pathway association analysis, the Reactome pathway database^43^ was used. We created a subset of the pathway database by selecting only the gene-sets containing at least one potential drug target and containing less than 300 genes. We also grouped the drugs in different classes according to their genomic target. In total we selected 94 pathways related to 57 drugs and drugs were classified into 11 classes. Pathway analysis was performed using the R Piano package (version 1.20.1)^47^. Circos plot was used to visualize the drug-pathway association^48^.

### Research Reproducibility

The source code of the Xeva package is open source under licence GPLv3 and available from GitHub (https://github.com/bhklab/Xeva). A complete software environment containing necessary data and code to reproduce the analysis and figures described in the manuscript is available at <u>Code Ocean</u>.

## RESULTS

### Xeva Follows PDX-MI Standards To Store Pharmacogenomic Data

We designed *XevaSet,* a new object class enabling integration of molecular and pharmacological profiles of PDXs (Supplementary Figure S1 and S2) following the recent Minimal Information for Patient-Derived Tumor Xenograft Models (PDX-MI) standard for reporting on the generation, quality assurance, and use of PDX models^32^. The PDX-MI standard ensures that all necessary clinical attributes of the tumor along with PDX-related essential experimental information, such as host mouse strain and passage information, is reported. Given that this information is crucial for downstream analysis and research reproducibility, we have implemented the *pdxModel* class (Supplementary Figure S2) which provides slots for PDX-MI variables. In this study, we curated the recent Novartis PDX Encyclopedia (PDXE)^28^ and created the PDXE *XevaSet* object (Figure 1) to investigate the consistency of gene expression patterns across passages and mine the pharmacogenomic data for known and new biomarkers predictive of drug response *in vivo*.

**Figure 1:**
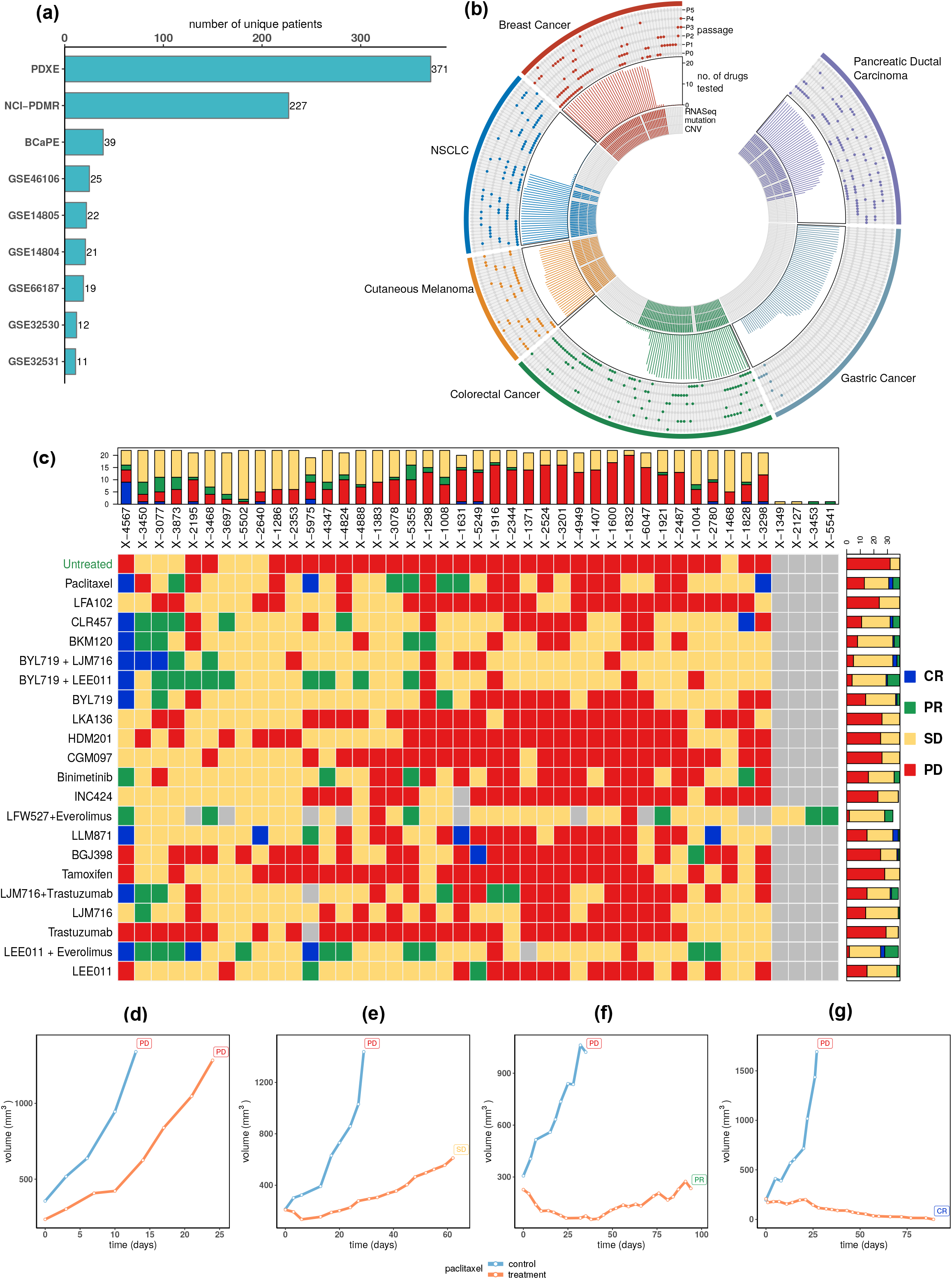
Xeva facilitates analysis of the PDX-based pharmacogenomic data. **(a)** Number of unique patients in different PDX based pharmacogenomic data-sets. PDXE refers to Gao et. al., 2015 data and GSE identifiers are from NCBI Gene Expression Omnibus (GEO) database. **(b)** Distribution of genomic, drug screening, passage related data and cancer types in PDXE data-set. For 350 patients belonging to 6 different cancer types, availability of genomic data (CNA, mutation and RNAseq) is shown in inner tracks. Number of drugs also includes untreated/control PDXs. Availability of passage specific gene expression data is shown in outer track. **(c)** Computation and visualization of response for PDX based drug screening in PDXE breast cancer data using Xeva mRECIST function. CR: complete response; PR: partial response; SD: stable disease; PD: progressive disease. Figure **(d), (e), (f)** and **(g)** shows control and treatment (paclitaxel) growth curve of PDXs for patient id X-2344, X-1004, X-3078 and X-5975, respectively. Visualization is done using *plotPDX* function in Xeva.

Xeva provides useful functions for analysis and visualization of PDX-based pharmacogenomic data. For every PDX model in the PDXE breast cancer dataset, mRECIST based response metrics was computed using the Xeva function *response* (Figure 1c). Heatmaps representing the mRECIST data cutaneous melanoma, colorectal cancer, gastric cancer, non-small-cell lung carcinoma (NSCLC) and pancreatic ductal adenocarcinoma can be found in Supplementary Figures S4 to S8, respectively. Visualization of PDX growth curves is an essential part of data quality control and analysis. Tumor growth curves for individual PDX models and for matched control-treatment models can be plotted using the *plotPDX* function. Examples of PDX tumor growth curves in control (untreated) and treatment (paclitaxel) conditions are shown in Figure 1d-g. All 4,483 tumor growth curves with matched control can be found in Supplementary Data File 1. PDX-related genomic data and linked drug screening response data can be extracted using the function *summarizeMolecularProfiles.* Similarly, the *drugSensitivitySig* function allows users to quantify the strength of each gene-drug association using a regression model.

### Gene Expression is Consistent Across Passages

PDX models are known to better represent the molecular characters of human tumor compared to simpler *in vitro* models^24,49–52^. PDXs show high similarity to patient samples for mutation rate^28, 53^ and CNA^24^ and methylation^54^ patterns, histological and molecular subtypes^24,50,55^. Before conducting drug screening, PDXs are passaged multiple times with the assumption that genomic characteristics of PDXs are stable across passages. Several studies have shown that mutation and copy number patterns are largely stable across passages^25,50,56–60^. However, a meta-analysis using inferred copy number data asserted that the genomic landscape of PDXs changes rapidly during passaging^61^. Given these contradictory results, it is vital to perform systematic analysis of non-inferred genomic data (i.e., gene expression profile) across passage^61^.

To address these concerns we sought to systematically analyze the changes in gene expression pattern across passages. We curated gene expression data for 661 PDX samples, derived from 371 patients and spanning from passage 0 (P0) to passage 5 (P5) (Figure 2a and supplementary Table S1). To visualize the consistency of gene expression patterns across passages, we performed a t-distributed stochastic neighbor embedding (t-SNE) analysis and projected the high dimensional PDX gene expression data into a two dimensional plot (Figure 2b). PDXs derived from a patient but belonging to different passages (defined as belonging to same lineage) were linked together in the t-SNE visualization. We observed that in the visualization PDXs from the same lineage are projected nearby even though coming from different passages. Next we computed the Pearson correlation coefficient for all pairs of PDX samples belonging to same lineage (Figure 2c). We found that the median Pearson correlation for related pairs is high (0.97) in comparison to non-related tissue specific PDX pairs (0.80) and the difference was statistically significant (p<0.0001). Collectively, these results strongly support that the gene expression profile of PDXs is consistent across different passages. For specific genes, however, the expression behaviour can vary across PDX passages, which may affect drug response depending on which passage is used for drug testing. We therefore assessed stability of each gene across passages by computing the intra-class correlation coefficient (ICC) for genes in samples from same PDX lineage for all tissue types and stratified by tissue type (Fig. 3). We found that the ICC values of genes in pairs of related PDXs are higher when compared to non-related pairs (Wilcoxon rank sum test p-value < 1E-15). Furthermore, the ICC values of genes are skewed towards high values indicating that the majority of genes have a stable expression pattern across PDX passages. Our results show that expression patterns of known biomarker genes such as *EGFR, ERBB2* and *MAP2K1* are stable across passage (Table 1 and Supplementary Data File 2). Consistent with Rubio-Viqueira *et al.*^52^ we found that expression of genes such as *VEGFA, MDM2* and *CDK4* is stable in pancreatic cancer PDXs.

**Figure 2:**
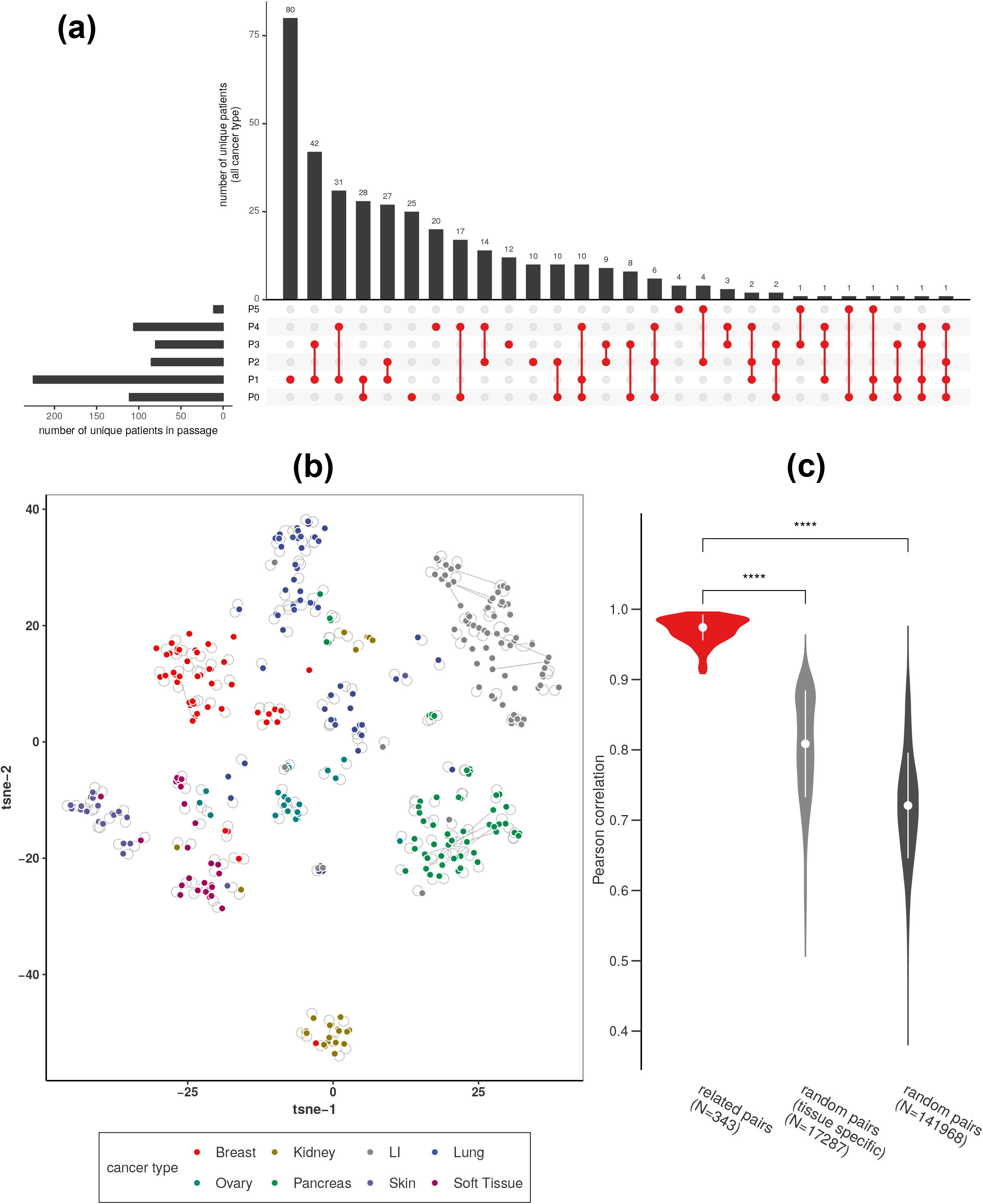
Gene expression landscape of PDXs is consistent across passages. **(a)** Distribution of samples in different passages. **(b)** TSNE analysis of gene expression data from different passages of the PDXs. Samples belonging to same lineage but belonging to different passages are linked together by line. **(c)** Pearson correlation for related sample pairs (belonging to same lineage) and randomly selected samples pairs. The correlation coefficient of related pairs is significantly higher then randomly selected pairs (p<0.001).

**Figure 3:**
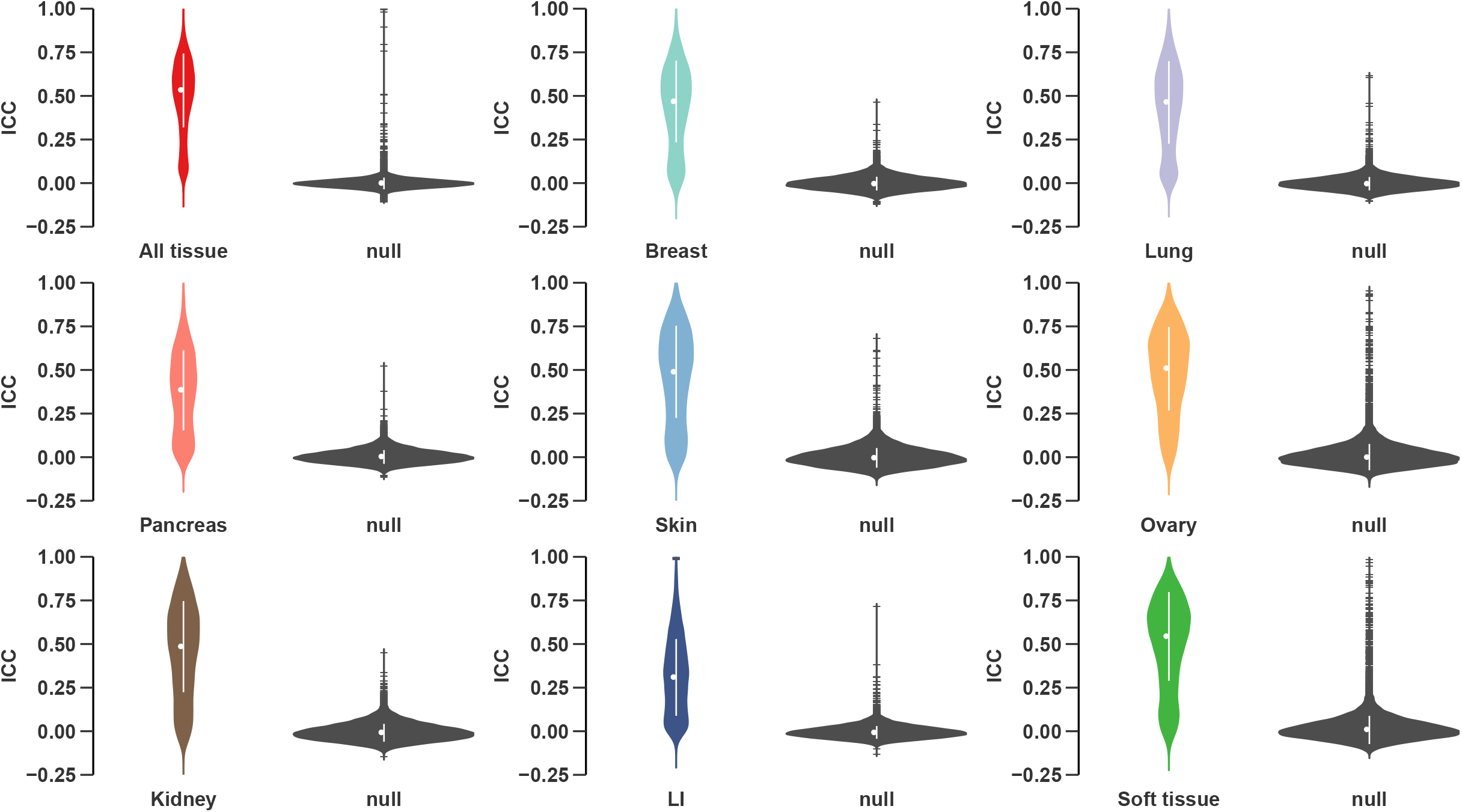
PDX maintains expression pattern of the genes across passages. Violin plot shows intraclass correlation (ICC) for genes across PDX passages for all samples and stratified by tissue type. Violin plot in black color represents ICC values for genes calculated using non-passage related (randomly selected) samples.

**Table 1:** Consistency of expression pattern of biomarkers across PDX passages. ICC values for biomarkers of FDA-approved anticancer agents in a complete dataset (all tissue) and stratified by tissue type. A higher ICC value indicates that expression pattern for the gene is consistent across passages. Biomarkers with ICC > 0.5 are considered preserved across passages and are highlighted in blue, or red otherwise.

| Biomarker | Drug(s) | ICC |  |  |  |  |  |
| --- | --- | --- | --- | --- | --- | --- | --- |
|  |  | All tissue | Breast | Lung | Pancreas | Skin | Ovary |
| EGFR | tyrosine-kinase inhibitor (erlotinib, gefitinib, saracatinib) | 0.74 | 0.64 | 0.70 | 0.50 | 0.42 | 0.78 |
| HGF | c-Met/HGF inhibitors (crizotinib) | 0.78 | 0.97 | 0.46 | 0.23 | 0.15 | 0.02 |
| MET | c-Met inhibitors (crizotinib, cabozantinib) | 0.72 | 0.51 | 0.63 | 0.41 | 0.79 | 0.84 |
| ERBB2 | HER2 inhibitors (lapatinib, trastuzumab) | 0.63 | 0.47 | 0.65 | 0.49 | 0.17 | 0.75 |
| ALK | ALK inhibitor (TAE684, crizotinib, ceritinib) | 0.61 | 0.42 | 0.74 | 0.07 | 0.25 | 0.76 |
| HGF | c-Met/HGF inhibitors (crizotinib) | 0.78 | 0.97 | 0.46 | 0.23 | 0.15 | 0.02 |
| MET | c-Met inhibitors (crizotinib, cabozantinib) | 0.72 | 0.51 | 0.63 | 0.41 | 0.79 | 0.84 |
| CDK4 | CDK4/6 inhibitor (palbociclib, abemaciclib) | 0.63 | 0.61 | 0.43 | 0.61 | 0.57 | 0.22 |
| CDK6 | CDK4/6 inhibitor (palbociclib, abemaciclib) | 0.67 | 0.75 | 0.64 | 0.57 | 0.63 | 0.73 |
| MAP2K1 | MEK inhibitor (binimetinib, trametinib, selumetinib) | 0.51 | 0.44 | 0.49 | 0.42 | 0.63 | 0.40 |
| MAP2K2 | MEK inhibitor (binimetinib, trametinib, selumetinib) | 0.42 | 0.38 | 0.35 | 0.42 | 0.59 | 0.50 |
| NQO1 | tanespimycin | 0.82 | 0.50 | 0.89 | 0.64 | 0.82 | 0.60 |
| MDM2 | Nutlin-3 | 0.68 | 0.72 | 0.38 | 0.70 | 0.64 | 0.62 |
| SRC | SRC inhibitors (saracatinib, dasatinib, bosutinib) | 0.49 | 0.31 | 0.20 | 0.17 | 0.41 | 0.67 |
| VEGFA | Bevacizumab | 0.46 | 0.30 | 0.48 | 0.44 | 0.43 | 0.07 |
| BRAF | Sorafenib | 0.48 | 0.50 | 0.51 | 0.48 | 0.62 | 0.29 |
| RAF1 | Sorafenib | 0.53 | 0.39 | 0.42 | 0.49 | 0.71 | 0.63 |
| FLT3 | Sorafenib | 0.21 | 0.27 | 0.01 | -0.01 | -0.03 | 0.34 |
| ABCC5 | 5-FU | 0.72 | 0.54 | 0.88 | 0.52 | 0.57 | 0.56 |
| DPYD | 5-FU | 0.60 | 0.58 | 0.63 | 0.68 | 0.48 | 0.54 |
| ERCC1 | cisplatin-based chemotherapy | 0.66 | 0.71 | 0.49 | 0.70 | 0.78 | 0.81 |

To identify pathways that are enriched with unstable genes, we performed a gene set enrichment analysis with the genome-wide ranking of genes based on their stability across passages (ICC values; Figure 4 and Supplementary File 3). We found that post-transcriptional mRNA processing pathways such as mRNA 3’end processing and mRNA splicing are enriched with unstable genes. It is well established that during proliferation and differentiation, cells adjust the mRNA and protein level by controlling the post-transcriptional mRNA processing pathways^62–64^ Therefore the instability of these post-transcriptional mRNA processing pathways might be attributed to the tumor growth in the PDX. Targeting these pathways using a drug in PDXs might result in inconsistent response at different passages.

**Figure 4:**
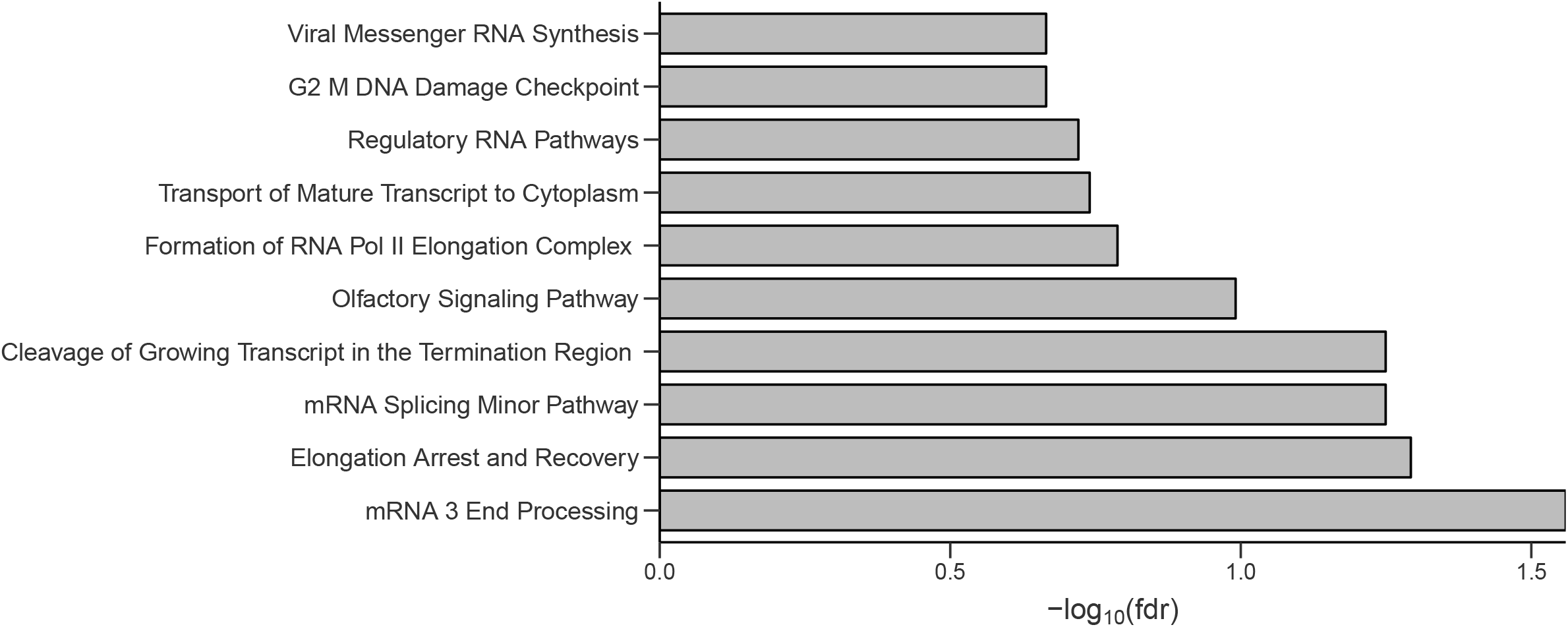
Pathways are stable across passages in PDXs. Barplot show top ten pathways with negative enrichment score in gene set enrichment analysis. Only one pathway has a statistically significant (FDR<0.05) negative enrichment score across PDX passages.

### *In Vivo* Biomarker Discovery

One of the main goals of pharmacogenomic studies is to find genomic biomarkers for drug response prediction. We evaluated the association between a molecular feature and response to a given drug (gene-drug association) in PDXE^28^ data. The PDXE data consists of a 1×1×1 experimental design where 60 compounds were tested across 277 PDXs. To model the tumor growth curve and to quantify the response of PDXs, we implemented several functions in Xeva. The Xeva function *plotPDX* provides an interface for the visualization of time versus tumor volume data of PDXs. Functions to compute drug response statistics include slope, area between curves, linear mixed effects model^39^, best average response and mRECIST^28^.

We employed Xeva to identify biomarkers for drug response prediction by computing gene-drug associations for drugs in the PDXE data and their corresponding known biomarker genes defined using OncoKB resource^46^. For the analysis, a PDX’s response to a drug is defined using best average response (BAR) and association was computed using concordance index. Analysis was done for each tissue type with gene expression, CNA and mutation data. *In vitro* drug testing has shown that the drug encorafenib can produce synergistic effects with binimetinib in cutaneous melanoma (CM)^28, 65^. This drug combination also shows synergistic effects in PDXs as 50% of tested PDXs show tumor shrinkage (Figure 5), while binimetinib and encorafenib monotherapy show tumor shrinkage in 40% and 25% of PDXs, respectively. For the drug combination of encorafenib and binimetinib, we found that NRAS mutation status is significantly associated with response (p-value=2.2E-05, Figure 5). Among PDXs with *NRAS* wild type status, 63.6% show a negative best average response to the binimetinib and encorafenib drug combination, while in the mutated category only 20% show a response.

**Figure 5:**
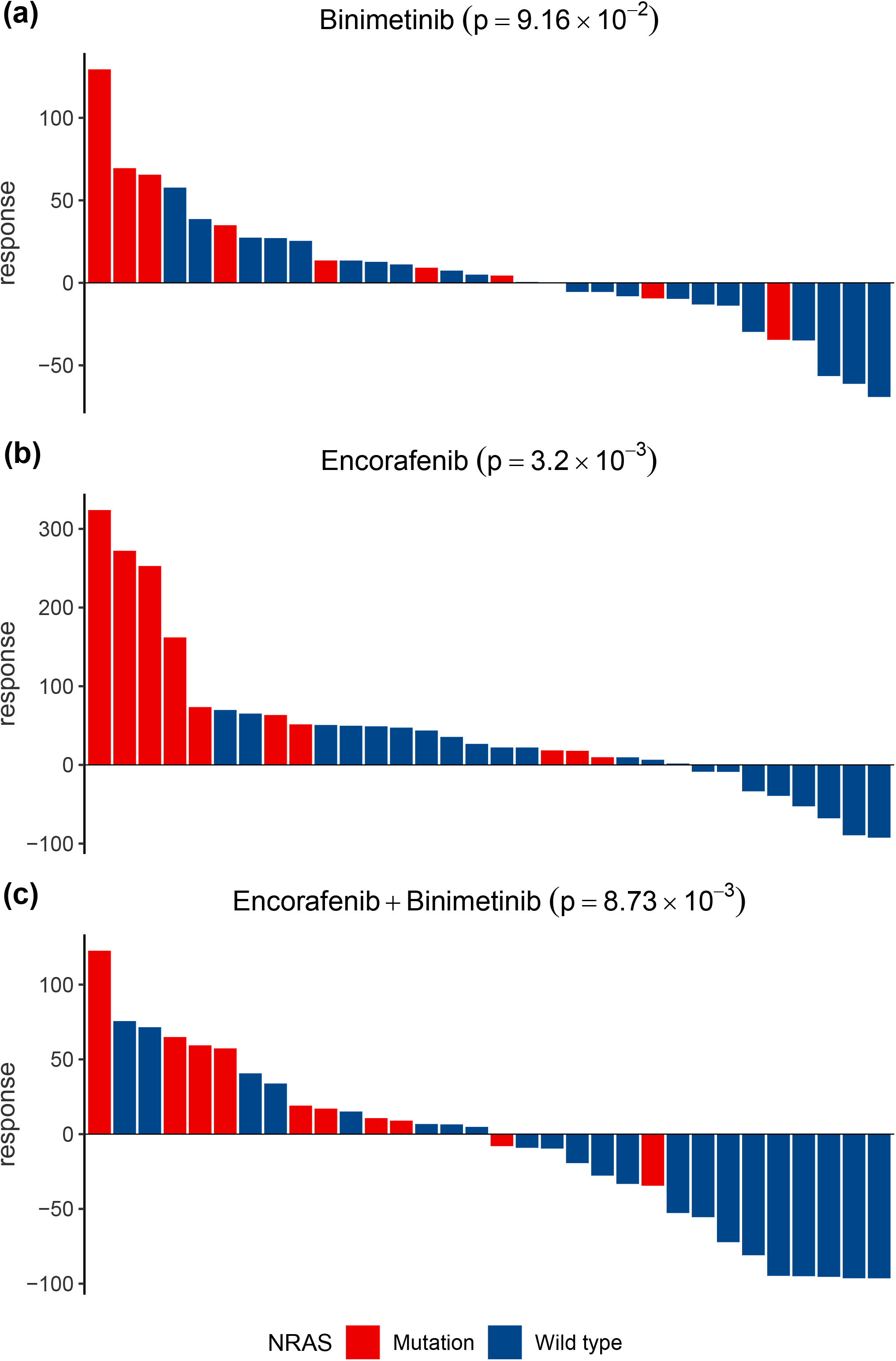
Xeva facilitates biomarker discovery and visualization. Waterfall plots show response of cutaneous melanoma PDXs for drug **(a)** Binimetinib **(b)** Encorafenib and **(c)** Binimetinib+Encorafenib. Each bar represent one PDX derived from a patient and color represent NRAS mutation status (red mutated and blue wild type). Response of the PDX is defined as best average response.

The drug trastuzumab (Herceptin) is a monoclonal antibody that targets the extracellular domain of the human epidermal growth factor receptor 2 protein (HER2) and inhibits the proliferation of tumour cells. In the breast cancer PDX data, we found that expression of the HER2 encoding gene *(ERBB2)* is significantly associated with trastuzumab response (CI=0.365; FDR=0.02, Figure 6). *ERBB3,* another member of the epidermal growth factor receptor (EGFR/ERBB) family, was also found to be associated with trastuzumab response (CI=0.36; FDR=0.026). *ERBB3* is known to be implicated in growth, proliferation, metastasis and drug resistance in tumors through interacting with *ERBB2*^66, 67^.

**Figure 6:**
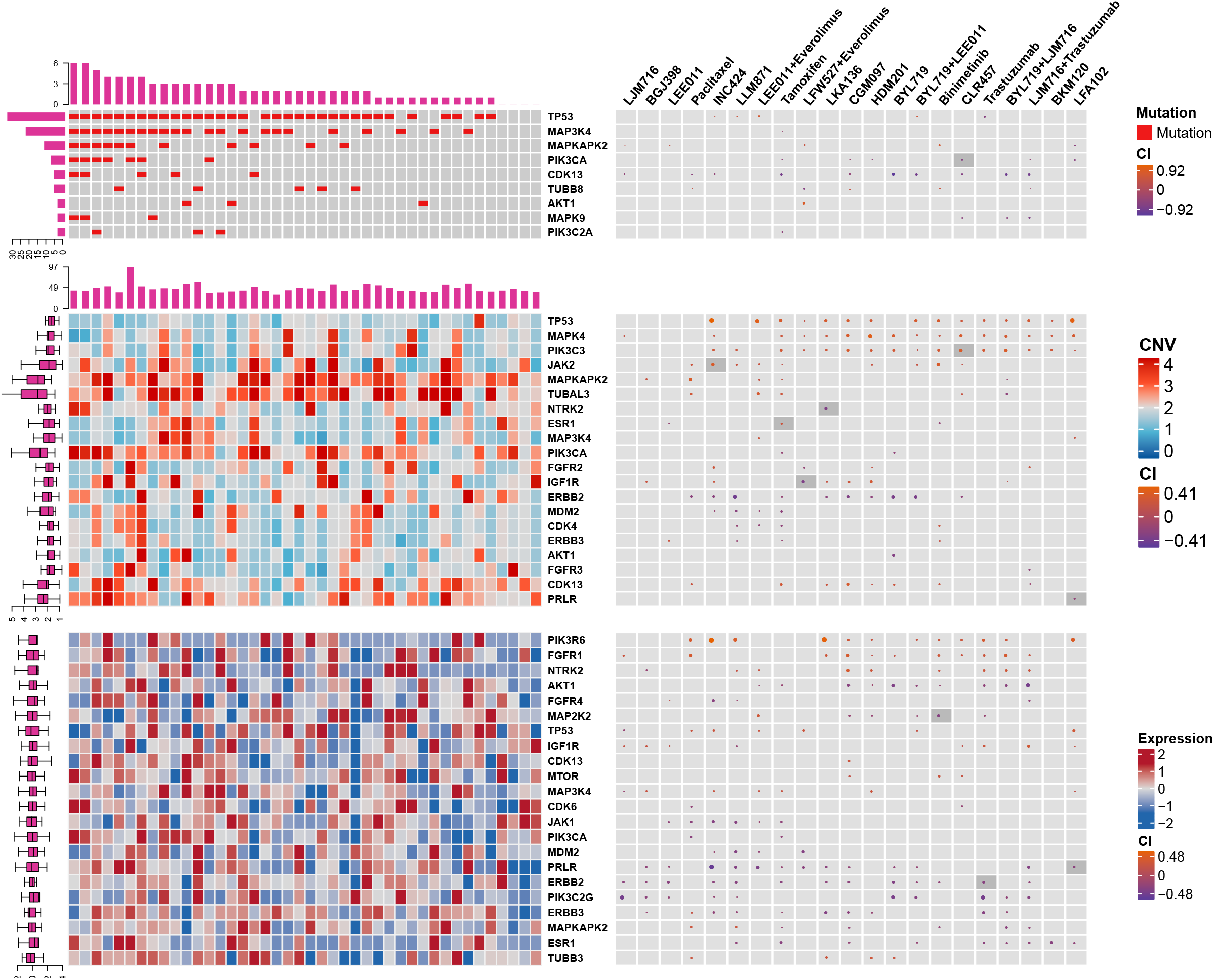
PDXs faithfully recapitulate known gene-drug associations. The left side of the figure shows mutation (top), CNA (middle) and expression (bottom) pattern of know biomarker genes. The right panel shows association (concordance index) between genomic features and response of the drug. For drugs the corresponding biomarker is highlighted using dark gray color if the association is statistically significant (FDR < 0.05).

The drug binimetinib is a targeted and potent mitogen-activated protein kinase kinase (MAP2K or MEK) inhibitor^68^. In breast cancer PDXs, expression of *MAP2K2* (CI=0.35; FDR=0.02, Figure-6) was significantly associated with binimetinib response. This gene belongs to the *MAP2K* kinase family and produces a protein which activates the MAPK/ERK pathway. Associations between potential biomarkers and PDX drug response for all tissue types can be found in the Supplementary Data File 4.

To gain understanding of the association between drug response and target pathway, we performed gene set enrichment analysis (Figure 7). Drugs were classified into 11 classes according to their known targets and the Reactome pathway database was used for gene set enrichment analysis. We found that EGFR signaling in cancer pathway is significantly enriched (FDR<0.05) in the EGFR class of drugs. Similarly, for MAPK class drugs, relevant pathways such as MAPK activation in TLR (Toll-Like Receptor) cascade and NFκB and MAPK activation by TLR are significantly enriched (FDR<0.05). Associations between each drug and reactome pathways can be found in Supplementary Data File 5.

**Figure 7:**
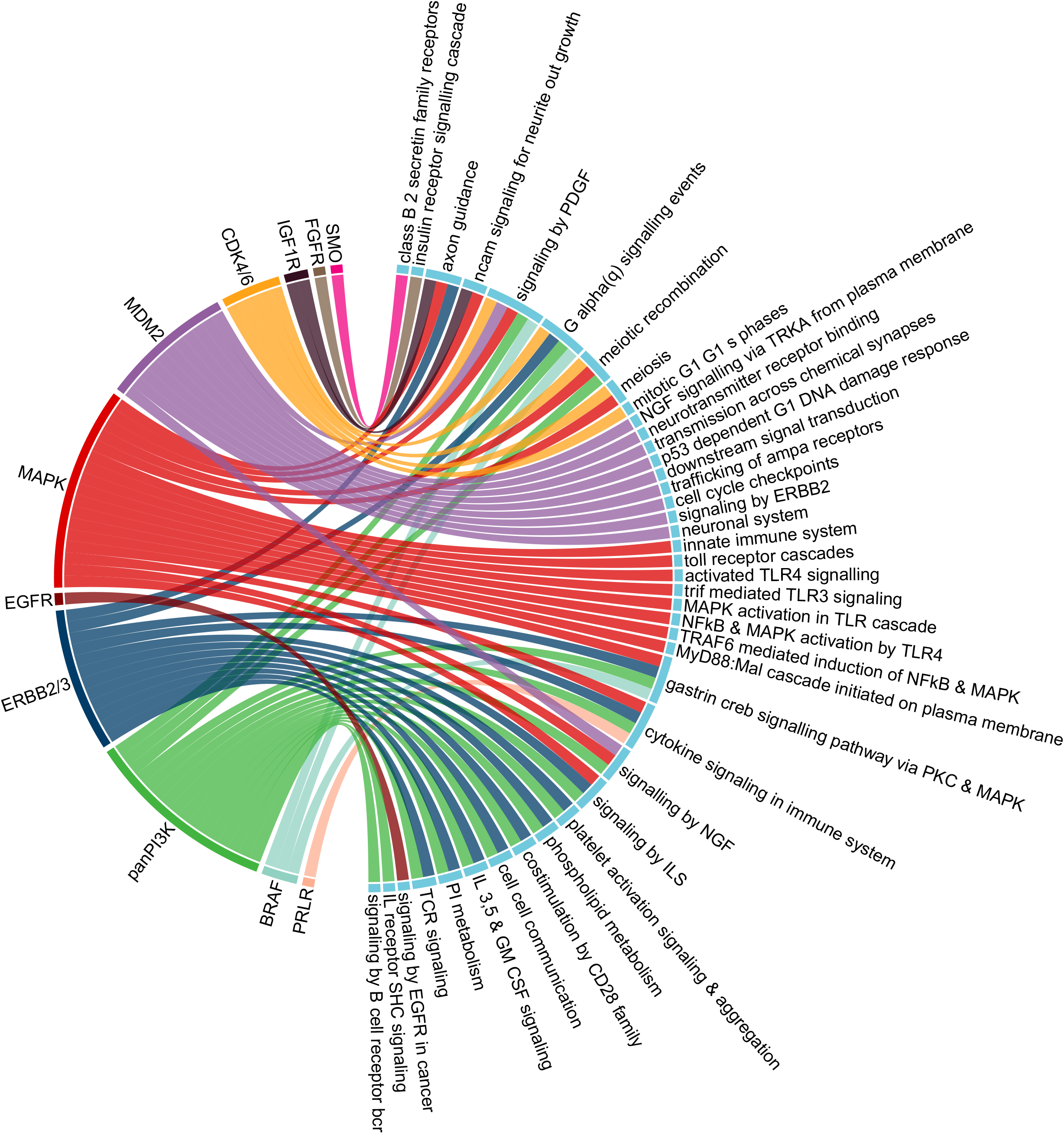
Pathways targeted by drugs are significantly enriched in PDXs. In the circos plot drug classes (left) and targeted pathways (right) are linked when the association is significant (gene set enrichment analysis FDR<0.05). Activity of EGFR class drugs have significant association with EGFR signaling pathway. Similarly MAPK class drugs show significant association with MAPK activation related pathways.

## DISCUSSION

Patient-derived xenografts are valuable models for cancer modeling and pharmacogenomic analysis. However, the translational potential of PDX preclinical models is highly dependent on the tools and techniques to processing and analyzing the data. Computational tools that enable standardized processing are thus an integral part of this line of research, and harmonized approaches can provide the community with accessible means for the analysis. The Xeva platform allows researchers to visualize and analyze the complex pharmacogenomic data generated during *in vivo* drug screening studies. The key strengths of the Xeva platform is its ability to store all metadata from a PDX experiment, link genomic data to corresponding PDX models and provide user friendly functions for analysis.

In a recent study, Ben-David *et. al.*^61^ analyzed changes in CNAs during PDX passaging using experimental and computational inferred copy number alterations (CNAs) from gene expression profiles. They concluded that the CNA landscape of PDXs changes rapidly with passage as within four passages 12.3% (median) of the genome was affected by model-acquired CNAs. While such an analysis is an important component of credentialing PDX as a pre-clinical platform, their analysis depended largely on inference rather than direct measurement of CNAs: for 84% (933 out of 1110) of PDX samples the copy number alterations were inferred from microarray-based gene expression data and lack matched normal tissue samples. As the authors stated, virtual karyotyping does not fully recapitulate the CNA observed from SNP microarrays or whole-exome sequencing. For the samples where DNA-based CNA profiles are available, the authors have reported a concordance of 0.82 between experimental (DNA based) and expression-based inferred CNA profile. However, directly assessing the gene expression pattern can provide better insight about changes in genomic landscape of PDXs across passages than the CNA profile. In this study, using Xeva, we have assessed the gene expression landscape of PDXs. Our results indicate that the gene expression landscape of PDXs is similar across different passages as the correlation between the related PDXs is very high. At the level of individual genes, we observe high consistency in expression patterns for the majority of genes. However caution is required when analyzing genes with low stability in PDX models. Lack of stability in gene expression may lead to inconsistent results when targeting proteins or pathways related to genes with low expression stability. We have provided a list of genes and their stability score which will help researchers to assess the consistency in expression patterns of genes of interest and thereby deciding if PDXs are suitable models for their (targeted) drug of interest or to study behaviour of a particular gene.

In our study, we found that the multiple known biomarkers predictive of drug response can be identified in the large Novartis PDXE dataset. Notably, the dataset used in this study has the 1×1×1 PDX experiment design (Supplementary Figure S9). Thus our results demonstrate that the 1×1×1 PDX experiment design is an adequate way to discover drug response predicting biomarkers at the population level. While the 1×1×1 experiment design is a cost- and animal-resource efficient way to find biomarkers, using Xeva, we have found cases where control (untreated) mice show a partial response (PR) or tumor shrinkage. Possible causes of such early tumor shrinkage might include experimental error, handling glitches or genetic properties of the tumor. In a 1×1×1 experiment design, lack of replicates makes it impossible to decipher the exact cause. Visualization and analysis of PDX response data using Xeva provides an efficient way to recognize such cases.

For biomarker discovery analysis, treatment response in PDXs is defined by the best average response (BAR)^28^. This metric of treatment response provides a continuous value, however it does not take into account the control arm of the PDX experiment. As PDX-based pharmacogenomics gains popularity, standardized metrics to define response are required, thereby taking into account the control arm of the PDX experiments. Such methods will also improve the biomarker discovery process. Xeva provides a standard tool for comparison of different PDX response metrics.

The Xeva package enables easy and efficient analysis of the PDX-based pharmacogenomic data. Xeva includes functions to link molecular features to drug response, therefore providing a unified framework for analysis and development of biomarkers of drug response.

## CONCLUSION

We developed the Xeva package to facilitate visualization, analysis and biomarker discovery in PDX pharmacogenomic data. We showed that PDX gene expression is consistent across passages. Our platform allowed us to confirm the existence of several drug response prediction biomarkers in a large PDX pharmacogenomic dataset. The reproducibility of known biomarkers and consistency in gene expression shows that PDX experiments are suitable for in vivo biomarker discovery or validation. Xeva is an open-source, flexible and timely tool in an era of increasing efforts to use PDXs as the main model system for cancer research. We envision Xeva will play a crucial role in PDX-based pharmacogenomic analysis, biomarker discovery and validation.

## ACKNOWLEDGEMENTS

ASM was supported by the Stand Up To Cancer Canada-Canadian Breast Cancer Foundation Breast Cancer Dream Team Research Funding, with supplemental support of the Ontario Institute for Cancer Research through funding provided by the Government of Ontario (Funding Award SU2C-AACR-DT-18-15). Stand Up To Cancer Canada is a program of the Entertainment Industry Foundation Canada. Research funding is administered by the American Association for Cancer Research International-Canada, the Scientific Partner of SU2C Canada. B.H.-K. was supported by the Gattuso Slaight Personalized Cancer Medicine Fund at Princess Margaret Cancer Centre, the Canadian Institute of Health Research, Natural Sciences and Engineering Research Council and The Terry Fox Research Institute fund. We like to thank the investigators of the Novartis Patient-Derived Xenograft Encyclopedia who have made their invaluable data available to the scientific community.

## FIGURES, TABLES and SUPPLEMENTARY INFORMATION

### Integrative Pharmacogenomics Analysis of Patient Derived Xenografts

**Figure S1:**
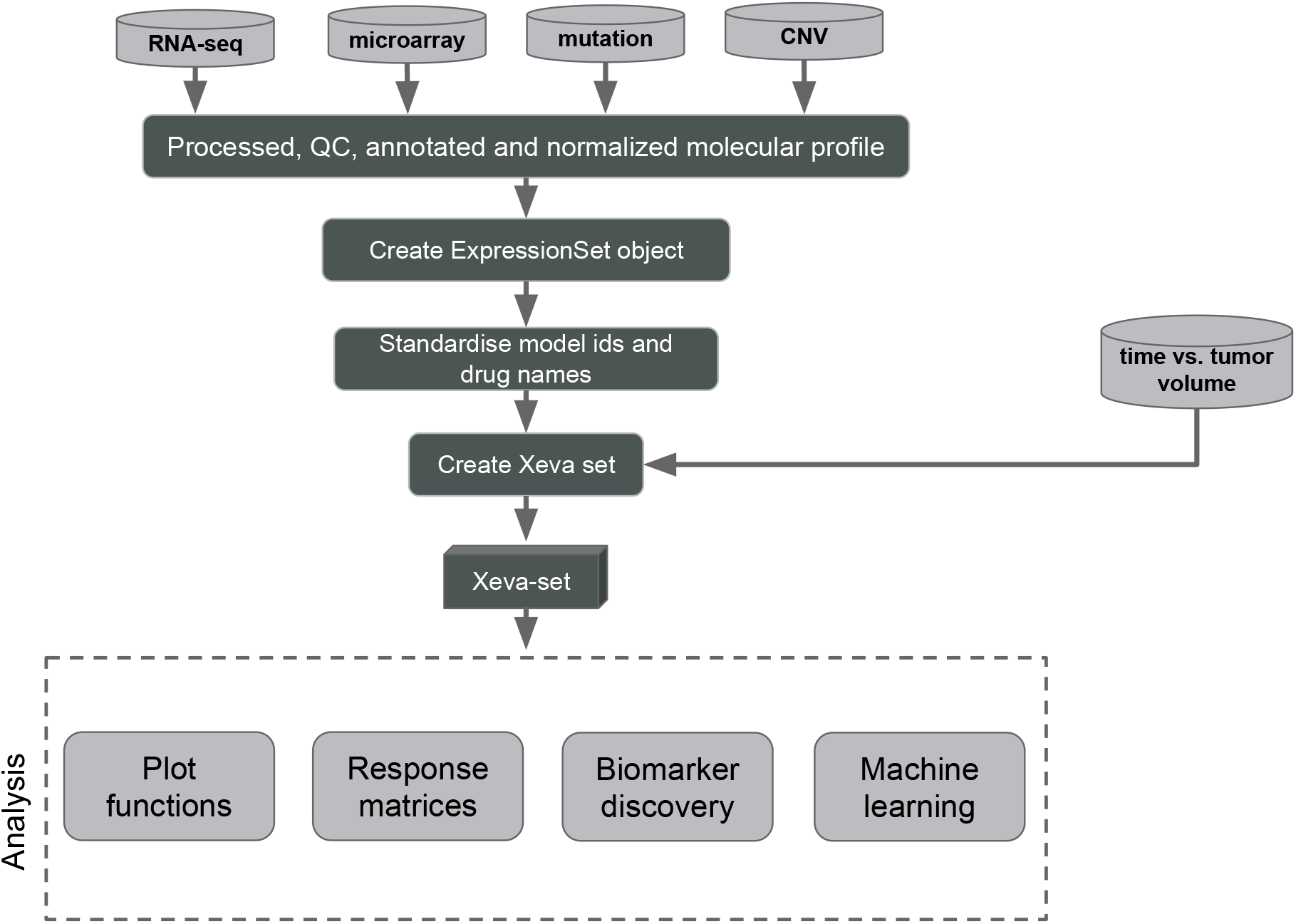
Description of the Xeva platform for PDX pharmacogenomics data management and analysis. Processed and curated genomic data (RNAseq, microarray, CNV and mutation) is formatted and stored along with drug response and experimental information in a unified manner. The Xeva tool provides several functions to analyze, summarize and plot the PDX data.

**Figure S2:**
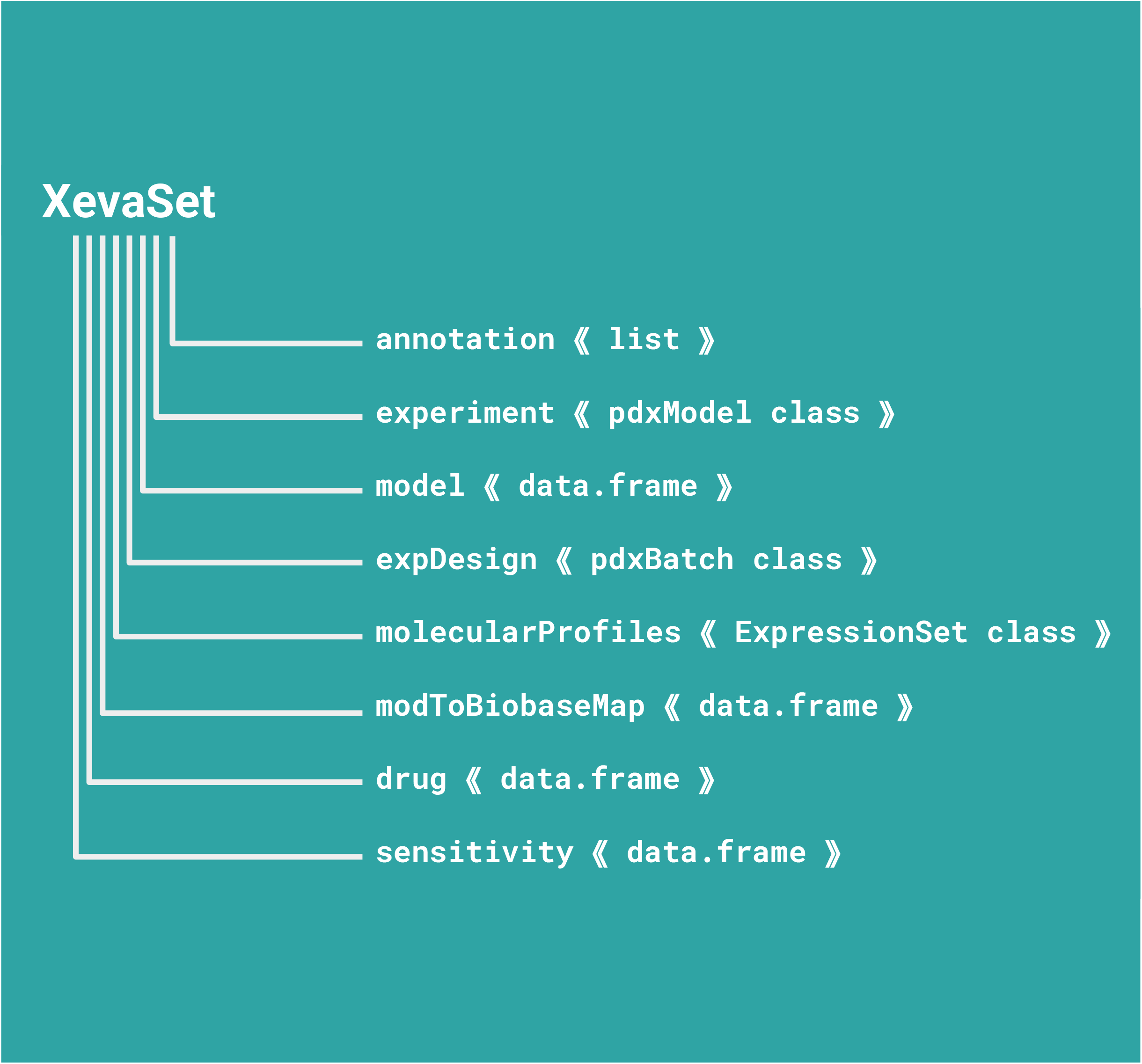
Schematic diagram of the XevaSet class. In Xeva package, XevaSet class combines different aspects of PDX based pharmacogenomic data. As an S4 class it provides slots for storing experimental data and associated molecular profiles. For each slot underlying class is represented in the bracket. For experiment slot, which holds time vs. tumor data of PDX, the detail schematic of underlying class pdxModel is shown in figure S3

**Figure S3:**
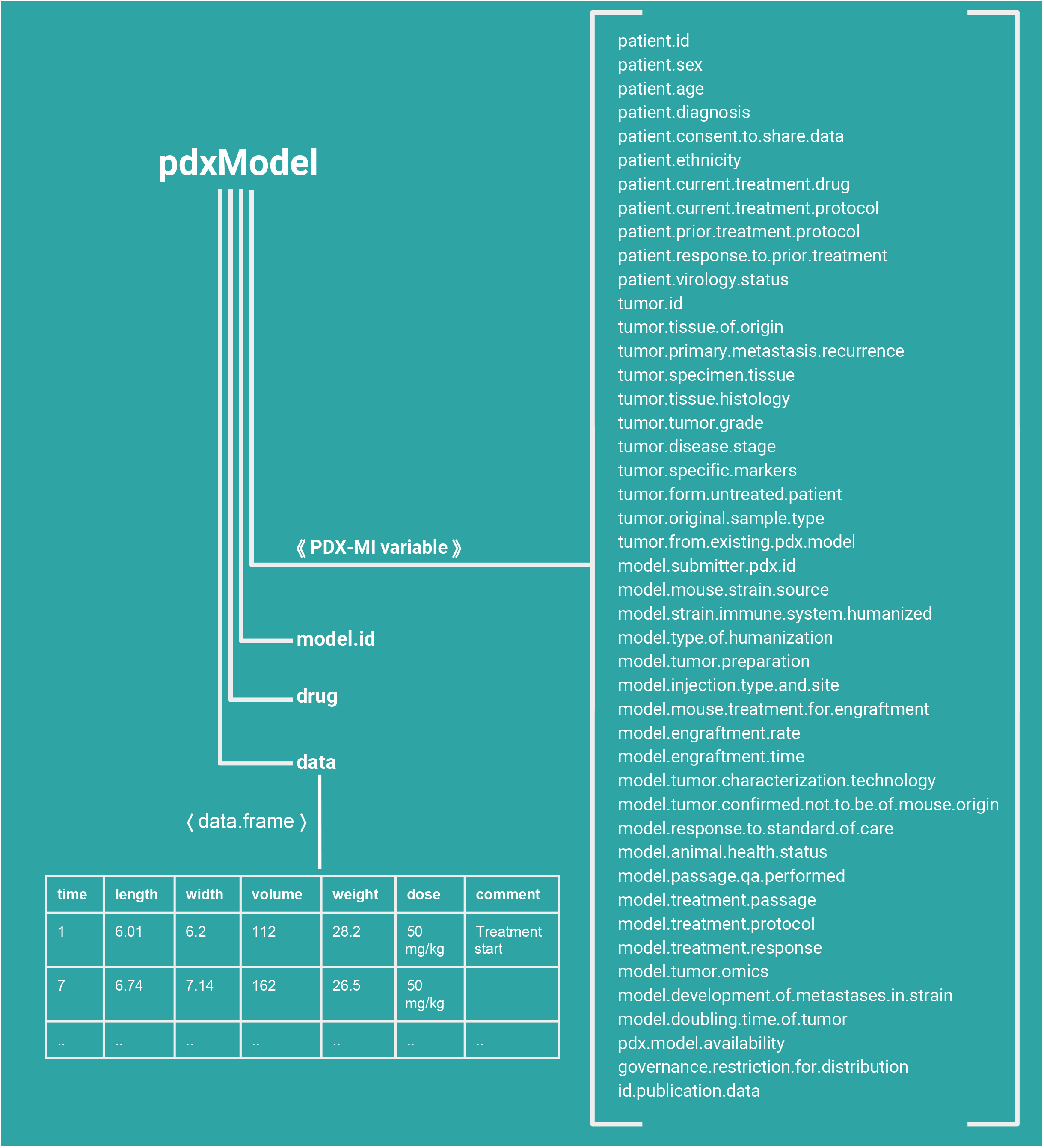
Schematic diagram of the pdxModel class in Xeva package. The pdxModel class is implemented as an S4 class which stores time vs. tumor volume data and associated meta data for an individual PDX model. It provides slots for different PDX-MI (PDX minimal information) variables. The data slot stores time vs. tumor volume information along with dose, mouse body weight and comments at individual time point.

**Figure S4:**
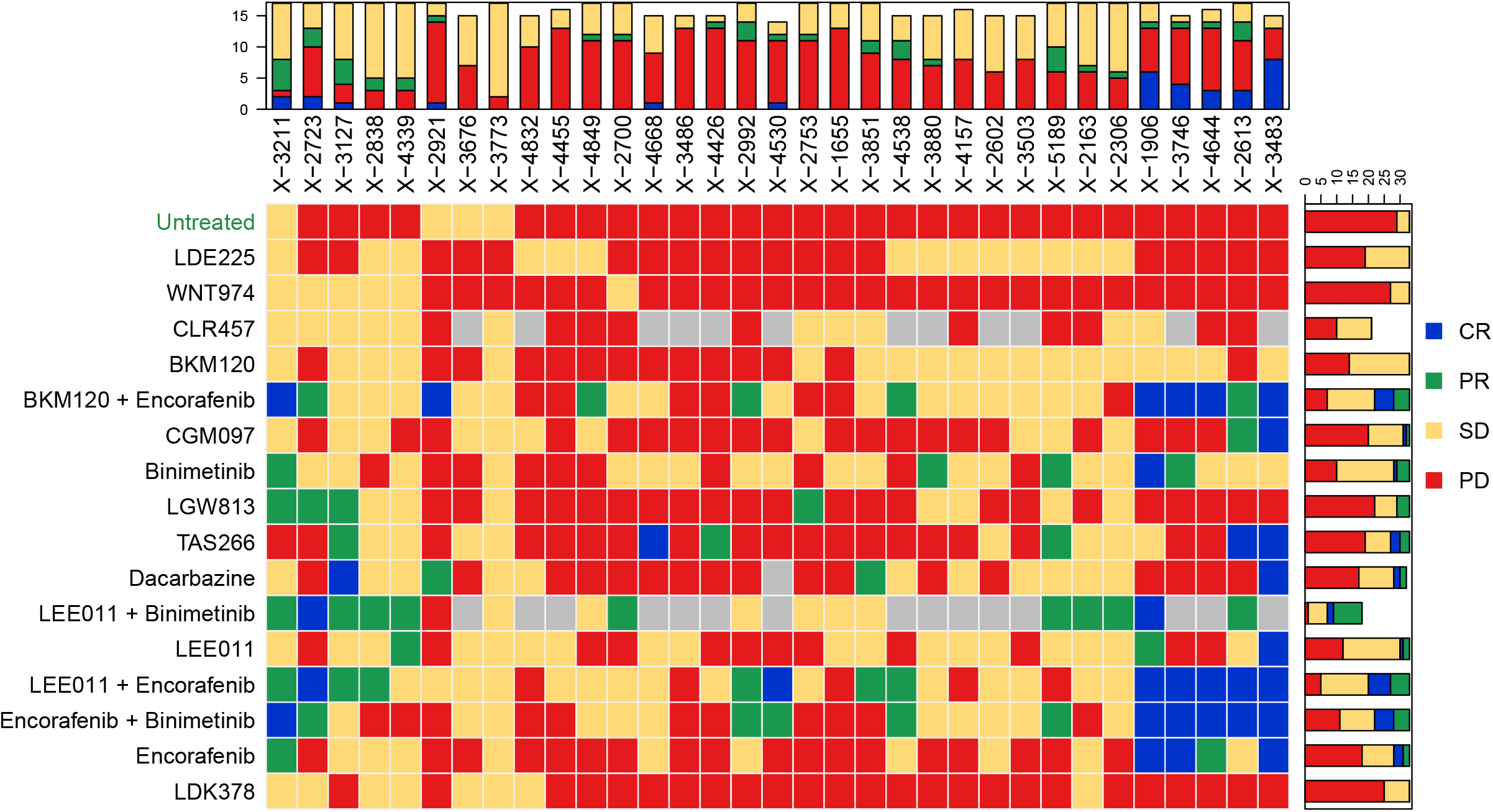
Computation and visualization of response for PDX based drug screening in PDXE cutaneous melanoma data using Xeva mRECIST function. CR: complete response; PR: partial response; SD: stable disease; PD: progressive disease.

**Figure S5:**
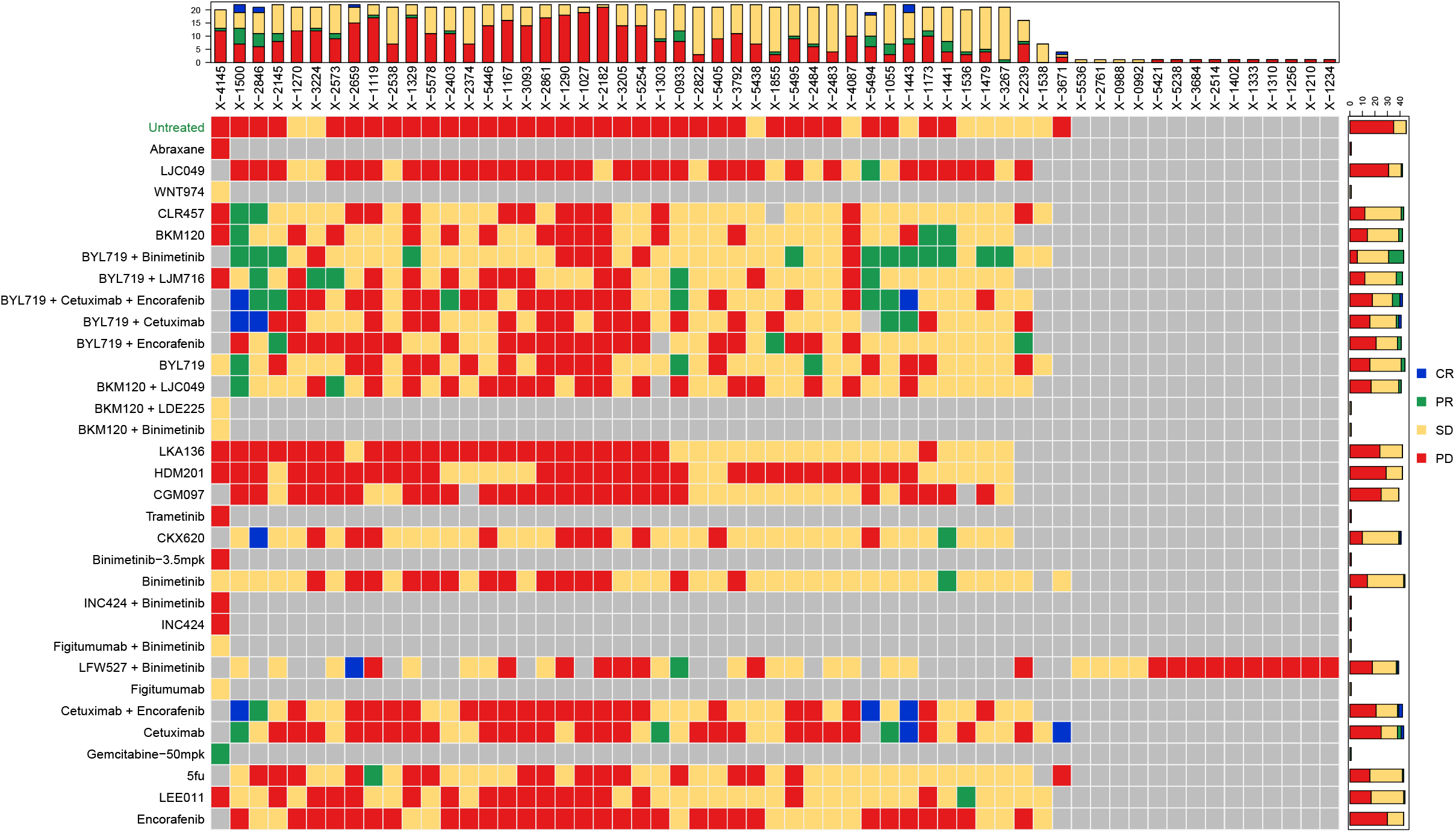
Computation and visualization of response for PDX based drug screening in PDXE colorectal cancer data using Xeva mRECIST function. CR: complete response; PR: partial response; SD: stable disease; PD: progressive disease.

**Figure S6:**
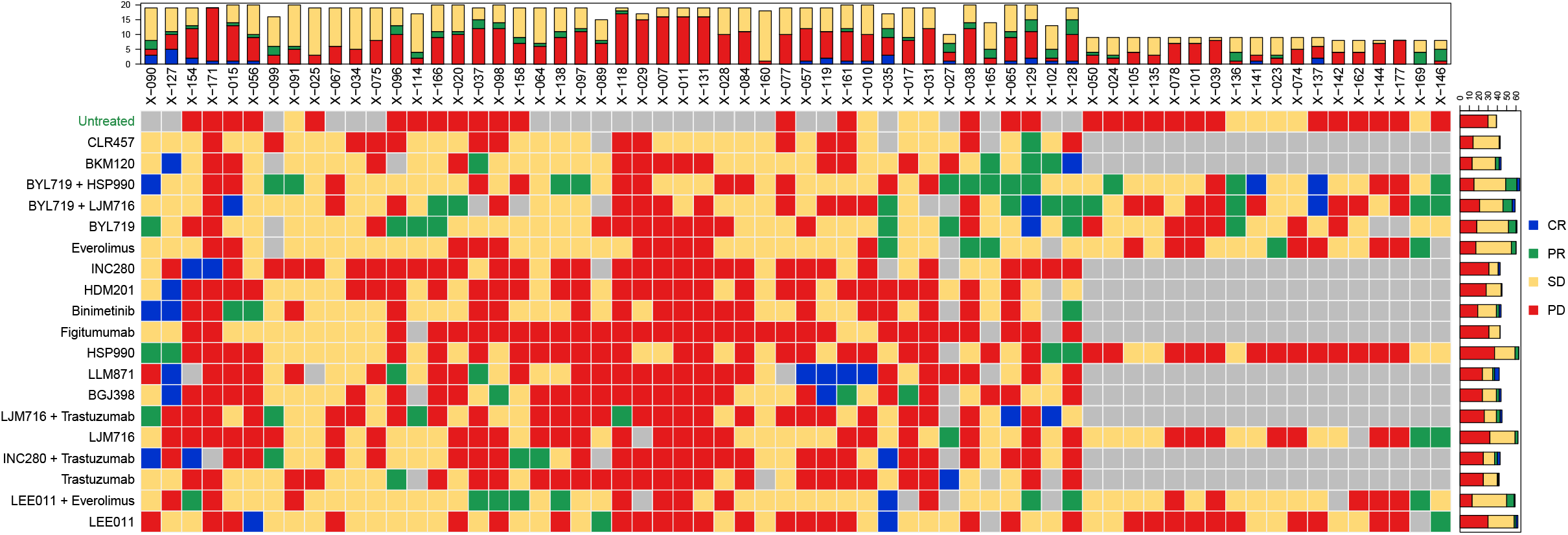
Computation and visualization of response for PDX based drug screening in PDXE gastric cancer data using Xeva mRECIST function. CR: complete response; PR: partial response; SD: stable disease; PD: progressive disease.

**Figure S7:**
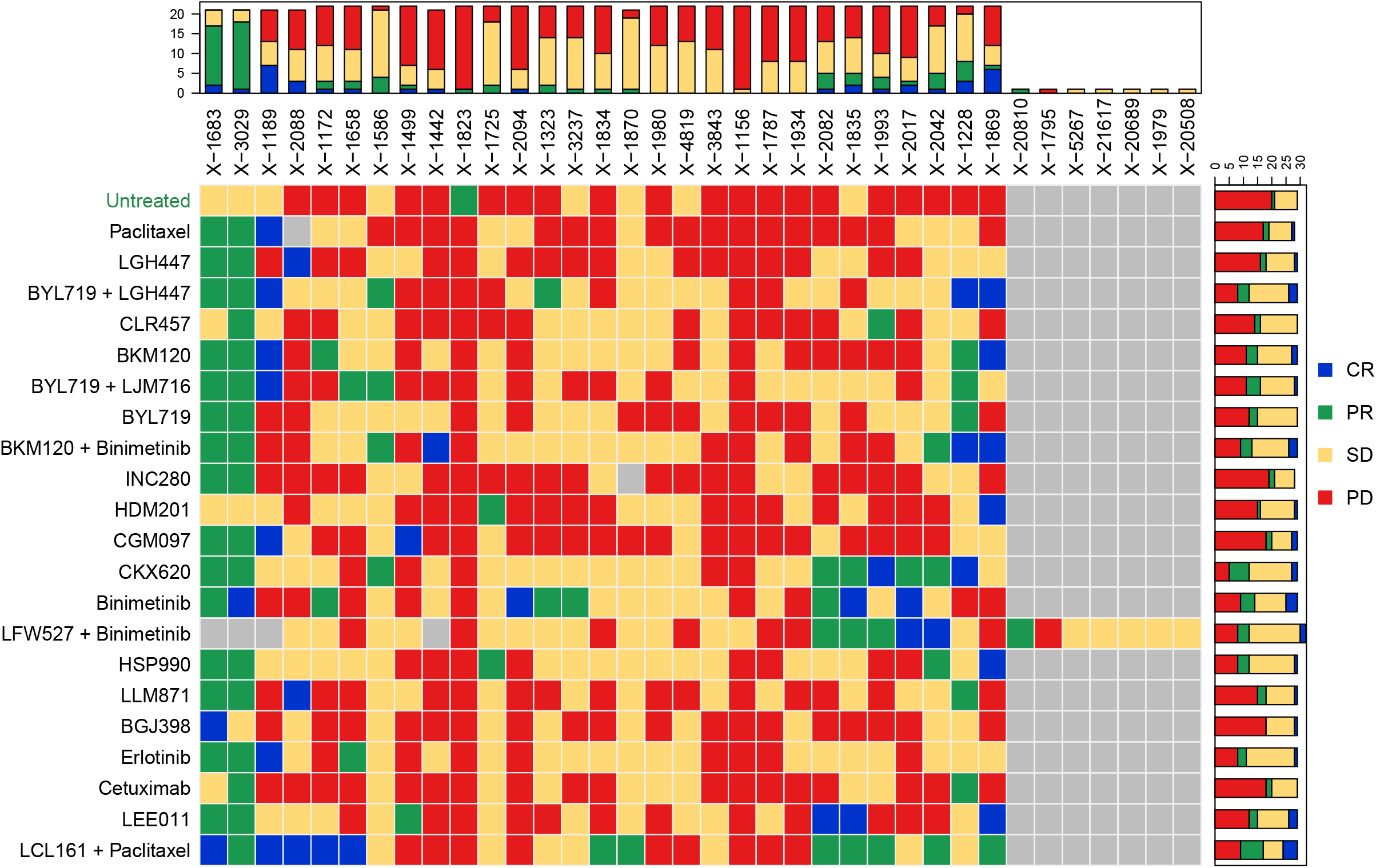
Computation and visualization of response for PDX based drug screening in PDXE non-small cell lung carcinoma data using Xeva mRECIST function. CR: complete response; PR: partial response; SD: stable disease; PD: progressive disease.

**Figure S8:**
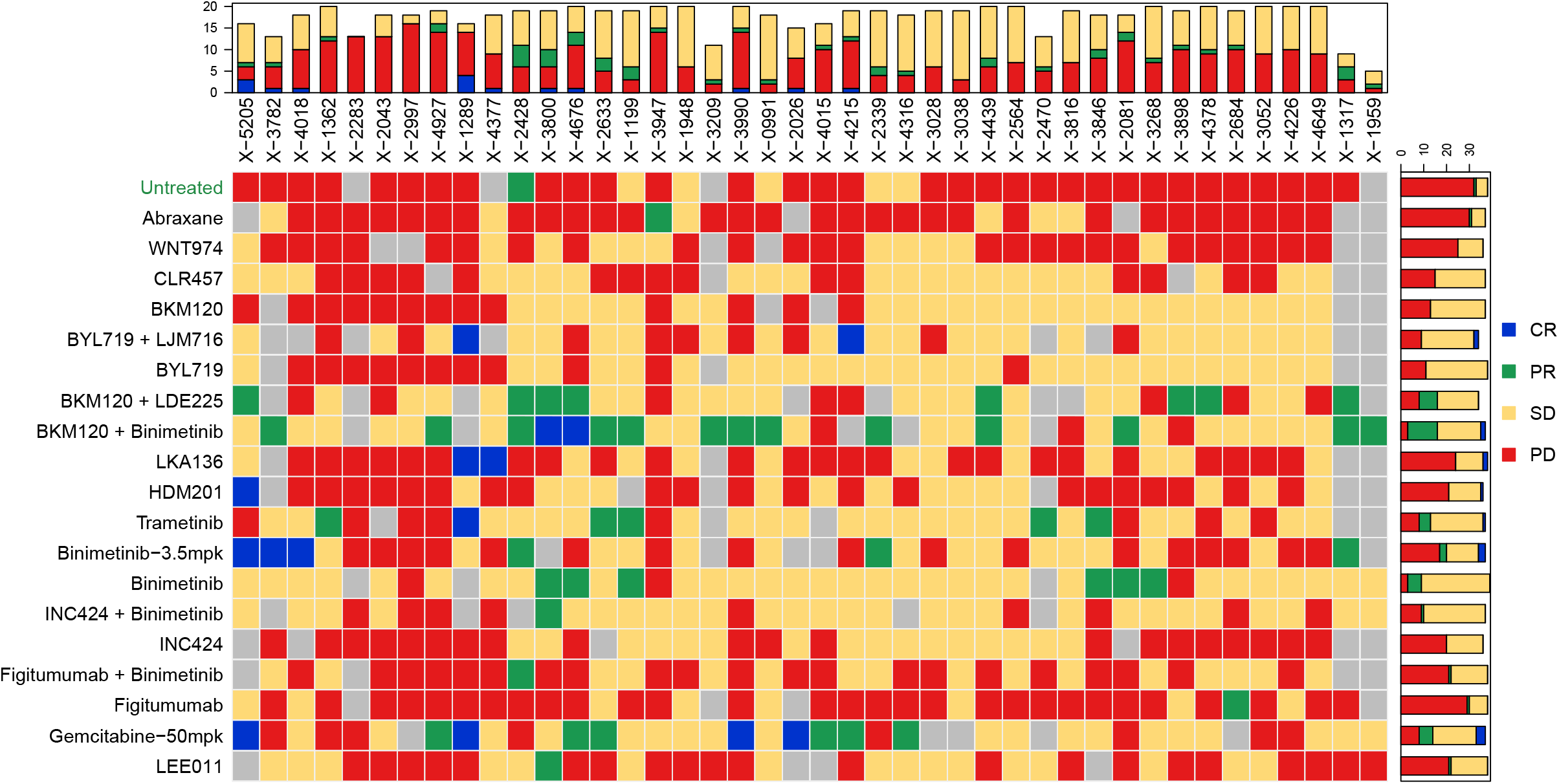
Computation and visualization of response for PDX based drug screening in PDXE pancreatic ductal adenocarcinoma data using Xeva mRECIST function. CR: complete response; PR: partial response; SD: stable disease; PD: progressive disease.

**Figure S9:**
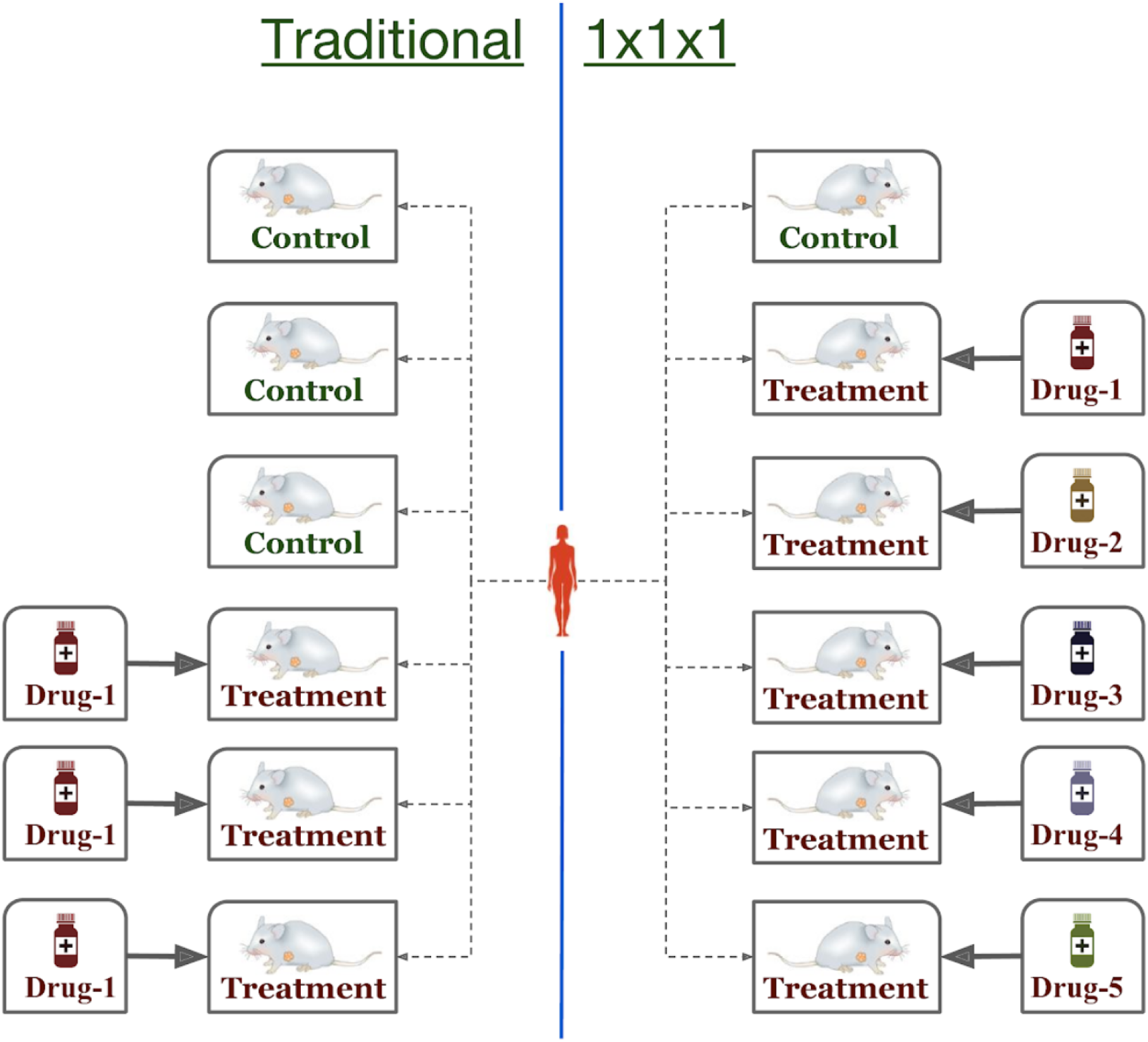
Different experimental design used in PDX based drug screening. In traditional design setting (left), multiple mouse models are created (replicates) for both control and treatment. In the 1×1×1 design only one control is generated for a patient and one for each treatment

**Table S1:**
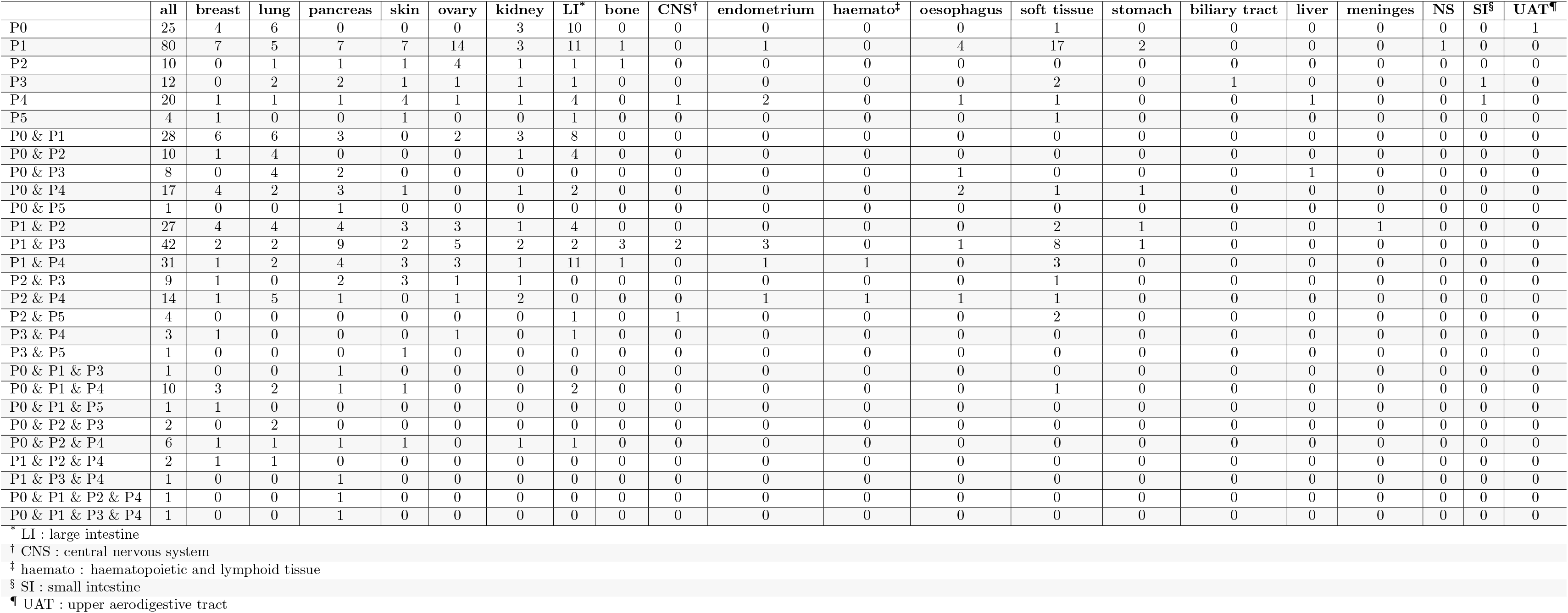
Number of unique patients in different passage in PDXE dataset, stratified by tissue type.

